# Dynamic integration of conceptual information during learning

**DOI:** 10.1101/280362

**Authors:** Marika C. Inhoff, Laura A. Libby, Takao Noguchi, Bradley C. Love, Charan Ranganath

## Abstract

The development and application of concepts is a critical component of cognition. Although concepts can be formed on the basis of simple perceptual or semantic features, conceptual representations can also capitalize on similarities across feature relationships. By representing these types of higher-order relationships, concepts can simplify the learning problem and facilitate decisions. Despite this, little is known about the neural mechanisms that support the construction and deployment of these kinds of higher-order concepts during learning. To address this question, we combined a carefully designed associative learning task with computational model-based functional magnetic resonance imaging (fMRI). Participants were scanned as they learned and made decisions about sixteen pairs of cues and associated outcomes. Associations were structured such that individual cues shared feature relationships, operationalized as shared patterns of cue pair-outcome associations. In order to capture the large number of possible conceptual representational structures that participants might employ and to evaluate how conceptual representations are used during learning, we leveraged a well-specified Bayesian computational model of category learning [1]. Behavioral and model-based results revealed that participants who displayed a tendency to link experiences in memory benefitted from faster learning rates, suggesting that the use of the conceptual structure in the task facilitated decisions about cue pair-outcome associations. Model-based fMRI analyses revealed that trial-by-trial integration of cue information into higher-order conceptual representations was supported by an anterior temporal (AT) network of regions previously implicated in representing complex conjunctions of features and meaning-based information.

## Introduction

One of the core functions of memory is the ability to use prior experience to optimize and facilitate decisions. A central challenge to this adaptive behavior, however, is the dense and continuous nature of experience. One approach to reducing this complexity is to make use of conceptual structure in the environment. Concepts can be formed on the basis of simple features (e.g. ‘has feathers’ to categorize animals as birds), however concepts are also thought to reflect broader forms of featural overlap, which can include the similarity of the relationships between features [1,2]. For example, concepts reflecting two different types of coins can be formed based on shared information about their value inside a fairground and at a convenience store. Tokens and medallions can be used pay for items at a fairground but are worthless at a convenience store, whereas quarters and dimes possess the opposite set of value relationships. Concepts like “carnival currency” or “world currency” can support decision-making and efficient learning about individual coins by allowing for inferences across different coins that share feature relationships [3].

Although recent neuroimaging investigations have begun to elucidate the brain regions that support conceptual or category membership on the basis of simple features [4–11], little is known about the neural mechanisms involved in the development and use of conceptual representations based on shared relationships across features. To address this question we combined a carefully designed associative learning task with computational model-based fMRI. Participants were scanned as they learned about sixteen pairs of novel cue objects and deterministically associated outcomes. Each trial began with the sequential presentation of a pair of object cues and a prompt to predict the associated outcome followed by response feedback. Critically, relationships between pairs of cues and outcomes formed a network of overlapping associations, where groups of cues shared identical cue pair and outcome associations, or identical feature relationships. These shared feature relationships could serve to simplify the learning problem from sixteen individual cue pair-outcome associations into four higher-order concepts, reflecting groups of cue-pair outcome associations containing individual cues with shared feature relationships (Fig 1). Importantly, this reduction of the learning problem was adaptive, and could allow for the acceleration of learning and facilitation of decisions in the task.

**Fig 1.**
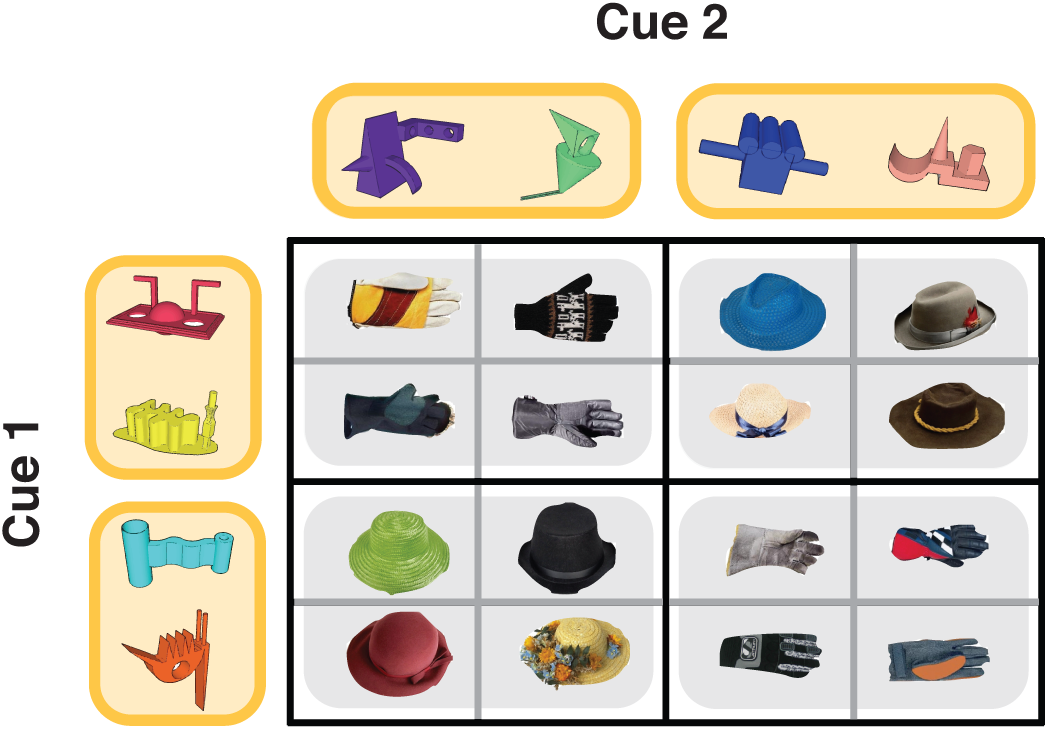
Task structure. Sixteen unique trial sequences of Cue 1, Cue 2, and outcome objects were constructed for each participant. In this example task structure, Cue 1 objects are presented along the y-axis, Cue 2 are objects presented along the x-axis, and associated Outcomes are presented in the center of the grid. For example, when the magenta Cue 1 is paired with the green Cue 2, the associated outcome is a glove. Individual cue objects each have a 50% chance of association with a Hat or Glove category outcome, requiring participants to use information about the Cue 1 – Cue 2 pair to make correct decisions. Cue 1 - Cue 2 - Outcome associations were fully crossed to create four pairs of cue objects that share feature relationships (highlighted in yellow). For example, both the magenta and yellow Cue 1 objects are associated with a glove category outcome when paired with the purple or green Cue 2 object and a hat category outcome when paired with the blue or tan Cue 2 object. This design gives rise to four groupings of Cue 1 - Cue 2 - Outcome associations where the corresponding cue objects create triplets with maximal conceptual overlap (highlighted in grey).

In order to elucidate the processes involved in building concepts based on shared feature relationships and to understand how they are represented in the brain, we turned to computational model-based fMRI. Specifically, we fit a well-specified Bayesian computational model of category learning [1] to trial-by-trial learning behavior, allowing for the generation of process-based estimates of dissociable aspects of the conceptual structure used by each participant during learning. We focused on two model estimates associated with the separate cue and outcome phases of each trial: a “Cue-based integration” parameter measuring the likelihood that a participant will incorporate cue pairs into an existing conceptual cluster, and a “Feedback-based updating” parameter, reflecting changes to the broader conceptual cluster space as participants receive and learn from response feedback.

Based on recent models positing the existence of two cortical networks that support memory-guided behaviors [12–14], we hypothesized that dissociable posterior-medial (PM) and anterior-temporal (AT) cortical networks would play key roles in representing Cue-based integration and Feedback-based updating. Specifically, a large number of investigations have linked regions in the AT network, including the perirhinal cortex (PRc) and orbitofrontal cortex (OFC), to the meaning of objects and integration of complex conjunctions of object features [4,15–23], suggesting that activity in the AT network might track Cue-based integration. On the other hand, PM network regions, including the parahippocampal cortex (PHc) [24–32], retrosplenial cortex (RSC) [33–36], and angular gyrus [37,38] have been shown to support memory for contextual information. Given that shared feature relationships in this task rely on the local context of each trial, or the trial-wise associations between cue pair and outcome, we might expect the PM network to also index Cue-based integration. We also hypothesized that the PM network would be preferentially involved in tracking Feedback-based updating, or trial-by-trial changes to the conceptual cluster space following response feedback, given proposals that that the PM network represents the full set of relevant relationships in the environment [13]. To assess whether parametric activity reflecting Cue-based integration and Feedback-based updating could be attributed to simple task accuracy, we also assessed PM and AT network activity during the cue and feedback periods of trials with correct outcome predictions relative to trials with incorrect outcome predictions. Finally, to validate PM and AT network-level grouping of individual brain regions, we also conducted an activation profile similarity analysis and tested whether regions in the same network displayed similar profiles of activation across the different experimental conditions.

## Materials and methods

### Subjects

Thirty-one (20 female) participants from the University of California at Davis community enrolled in the experiment. Two participants were excluded due to falling asleep inside the scanner, one participant was excluded due to excessive motion, and three participants were excluded due to issues with scanner protocol specifications. Of the remaining 25 participants (17 female), all had normal or corrected-to-normal vision, were native English speakers, and were 18 to 31 years of age. Informed consent was obtained in a manner approved by the Institutional Review Board at the University of California at Davis. Participants were paid $50 for their participation, and received additional compensation for the proportion of responses made above chance level on their best learning run (maximum additional payment of $5).

### Stimuli

To control for any use of semantic or perceptual information in learning cue pair-outcome associations, eight novel object stimuli were manually generated using Google SketchUp software (http://www.sketchup.com). Cue objects were designed to be visually distinctive in shape and color. Eight unique hat and eight unique glove outcome objects were selected from a stimulus database of objects [39].

### Design

Participants were scanned while completing six runs of a learning task. Each trial of the learning task began with the sequential presentation of two cue objects (Cue 1 and Cue 2) and an associated outcome. Importantly, each individual Cue object was associated with a 50% probability of predicting a hat or glove category outcome, requiring participants to integrate the combination of Cue 1 and Cue 2 to correctly predict the associated outcome category (Fig 1). Cue pair-outcome associations were generated by randomly assigning four of the eight novel objects to Cue 1, and the remaining four novel objects to Cue 2. To create a higher-order conceptual structure, cue pair-outcome associations were crossed to create pairs of individual Cue objects that shared feature relationships, or that shared patterns of cue pair-outcome associations (Fig 1, cues that share feature relationships highlighted in yellow). We reasoned that cue-pair outcome associations comprised of Cue 1 and Cue 2 objects with shared feature relationships would have maximal conceptual overlap (Fig 1, highlighted in grey).

### Experimental procedure

The experiment was comprised of four parts: unscanned target detection practice, one scanned pre-learning target detection run, six runs of scanned learning, one scanned post-learning target detection run, and a final unscanned memory test. Only the data from the scanned learning period were included for analysis.

During the six scanned learning runs, participants were presented with trials consisting of sequentially presented Cue1 – Cue 2 – outcome associations (Fig 2). Each trial began with the presentation of a Cue 1 object in the center of the screen for 2 seconds, followed by a blank screen for 500 ms. After the presentation of Cue 1, a Cue 2 object was presented for 2 seconds with the words ‘hat’ and ‘glove’ printed underneath, referring to the possible category-level outcomes. Participants were asked to use their pointer or middle finger to predict the category-level outcome associated with the current Cue 1 – Cue 2 pair.

**Fig 2.**
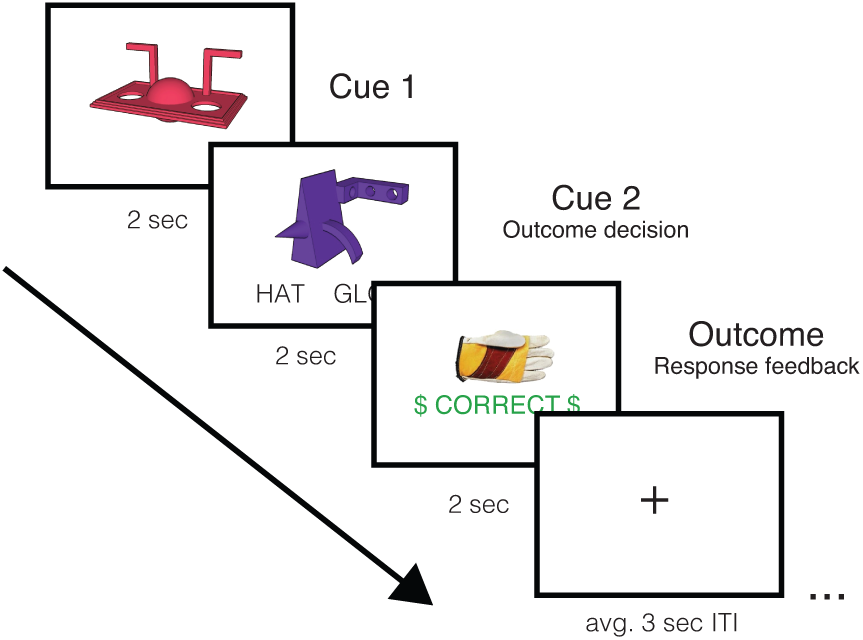
Trial sequence during scanned learning. Participants were presented with Cue 1, Cue 2 and Outcome feedback information sequentially.

Response buttons were counterbalanced across participants. Following the 2 second Cue 2 period, a 500 ms blank screen was presented, followed by outcome information and response feedback. Specifically, participants were presented with a unique hat or glove associated with the Cue 1 – Cue 2 pair, as well as feedback on the category-level decision made during the Cue 2 period. Each Cue 1 – Cue 2 – Outcome association was presented three times per run, and trial order was pseudo-randomly determined by drawing without replacement from the sixteen Cue 1 – Cue 2 – Outcome associations three times, with no back-to-back repetitions of individual associations. This resulted in three iterations through the full set of sixteen cue pair-outcome associations, for a total of forty-eight trials per learning run. A variable ITI with a static fixation cross followed each trial, and lasted between 1 and 4 seconds, with a mean of 3 seconds. Each run lasted 8 minutes and 8 seconds. In addition to trial-by-trial feedback, participants were presented with information about the proportion of trials they had answered correctly at the end of each learning run.

## Model-based analyses

Although the learning task was comprised of sixteen individual cue pair-outcome associations, the contingencies between cue objects and outcomes were designed such that participants could facilitate learning by integrating across cue pairs that contained Cue items with similar cue pair-outcome relationships (Fig 1). However, we expected large individual differences in the degree to which a participant could learn and use the conceptual structure of the task to guide learning. To accommodate this variance and to gain leverage on the processes that supported learning, we used the Rational Model of Categorization (RMC) to model behavioral data from the learning task [1]. We chose to apply the RMC based on previous theory [40] linking clustering mechanisms to key regions of interest and related model-based fMRI studies [5,41].

The RMC assumes that categories are learned by clustering similar stimuli together. Suppose a learner has observed *n* - 1 stimuli {*x*_*1*_*x*_*2*_…, *x*_*n-1*_} with corresponding category labels {*y*_*1*_, *y*_*2*,_ …, *y*_*n-1*_}. Each stimulus is fit into a cluster {*□*_*1*_, *z*_*2*_…, *z*_*n-1*_}. In the context of present study, *x*_*i*_ is a pair of cues presented at the *i*th trial, and *y*_*i*_ is a corresponding category outcome. An exact object which followed a cue pair (e.g., green hat and black glove) is denoted as 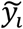 If the cue pair *x*_*i*_ was fit into the *j*th cluster, *z*_*i*_ equals to *j*.

Now, let us suppose *w* (0 < *w* < *n*) clusters have been formed after *n* - 1 trials. Then, the probability that the cue pair at the *n*th trial is judged to be from category *h* follows Bayesian inference:

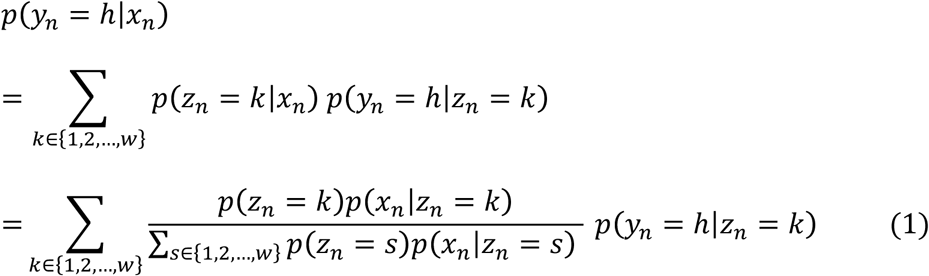

The three terms in Equation 1 is described below in turn.

First, the probability that the *n*th cue pair fits into the *k*th cluster is given by

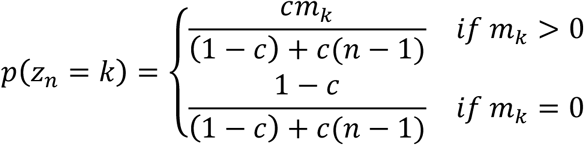

where *c* is a parameter called the coupling probability, and *m*_*k*_ is the number of cue pairs already assigned to the *k*th cluster.

The coupling probability is a single value for each participant that reflects the sensitivity in generating new clusters or to linking information to existing clusters in memory. A smaller coupling probability, for example, indicates that a new cluster is more likely to be created to accommodate the *n*th cue pair. Conversely, a larger coupling probability indicates that information is likely to be linked with an existing cluster. Thus, individual differences in learning can be captured by allowing the coupling probability parameter to vary across participants.

We assume that cues are independent of each other [1,42]:

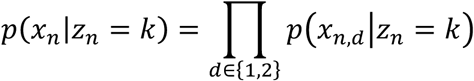

Here, *x*_*n,d*_ denotes the *d*th cue in the cue pair at the *n*th trial. This term is calculated with

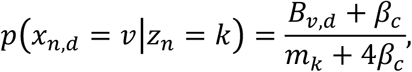

where *β*_*c*_ is the sensitivity parameter for a cue, and *B*_*v,d*_ the number of cue pairs in the *k*th cluster whose *d*th cue is *v*.

Similarly, the probability that the *n*th cue pair is from category *h* given a cluster is given by

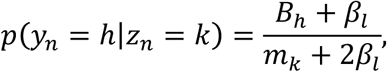

where *β*_*l*_ is the sensitivity parameter for a category, and *B*_*h*_ is the number of cue pairs in the *k*th cluster whose category is *h*. Unlike the coupling probability, the sensitivity parameter was not allowed to vary between participants.

After observing outcomes associated with each cue pair, a learner assigns a cue pair to a cluster. This cluster assignment also follows Bayesian inference, where the probability that the *n*th cue pair fits into the *k*th cluster is given by

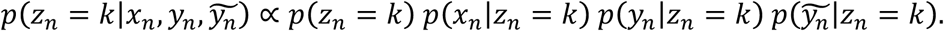

The last term is given by

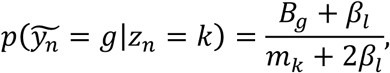

where *β*_*g*_ is the number of cue pairs in the *k*th cluster which is associated with object *g* (e.g., green hat).

This learning by clustering is probabilistic, and the same parameter values can result in different cluster formulations. To account for this stochasticity, we simulated the model 2,000 times with one particle when evaluating a set of parameters [42]. This stochastic learning allows for a characterization of dissociable processes involved in using the conceptual cluster space.

To obtain trial-by-trial measures, we estimated the maximum a posteriori of parameter values (the coupling probability and the sensitivity parameters) using the Bayesian optimization framework. The prior distribution for the coupling probability was the uniform distribution between 0 and 1, and the prior distribution for the sensitivity parameters was the uniform distribution between 0.01 and 10. The estimated parameter values are: the coupling probability ranges from 0.0002 to 0.0373 with a mean of 0.0088, and the sensitivity parameters are 0.01 for both cue and outcome category. With these parameter values, we took mean average of trial-by-trial measures from the 2,000 simulations.

In order to elucidate the involvement of PM and AT networks, we focused on two model-derived measures reflecting the trial-by-trial development and use of concepts during different phases of each trial: “Cue-based integration,” and “Feedback-based updating.” Cue-based integration is a measure assessing the likelihood that participants will assign or integrate a pair of cues to an existing conceptual cluster rather than generating a novel conceptual cluster for the cue pair. Using the above notation, the Cue-based integration estimate is given by 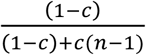. Feedback-based updating, on the other hand, assesses how much the conceptual cluster space changes following feedback. Thus, Feedback-based updating indicates the extent to which one learns and modifies their knowledge based on the outcome of each individual trial. Feedback-based updating is quantified as the Kullback-Leibler divergence between the probability distributions over the clusters before and after the feedback: *p* (*z*_*i*_|*x*_*i*_) and 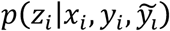.

## FMRI methods

MRI scans were acquired at the UC Davis Imaging Research Center using a 3T Siemens Skyra equipped with a 32-channel head coil. Participants were supplied with earplugs to attenuate scanner noise, and head padding was used to reduce motion. Stimuli were presented visually on a screen at the back of the scanner, and viewed through a mirror attached to the head coil. T1-weighted structural images were acquired with a magnetization-prepared rapid acquisition gradient echo (MPRAGE) pulse sequence (1 mm^3^ voxels; matrix size=256 × 256; 208 slices) and images sensitive to BOLD contrast were acquired using a whole-brain multiband gradient echo planar imaging sequence (3 mm^3^ voxels; TR = 1220ms; TE = 24ms; FA = 67°; multiband acceleration factor = 2; 38 interleaved slices; FOV = 192 mm; matrix size = 64 × 64).

Preprocessing and analysis of functional MRI data was conducted with FSL (FMRIB Software Library, https://fsl.fmrib.ox.ac.uk/fsl/fslwiki). The first 7 volumes of each functional run were discarded to allow for signal normalization. EPI volumes underwent motion correction using MCFLIRT [43], and were also highpass filtered (0.01 Hz) and spatially smoothed (6 mm full-width at half maximum Gaussian kernel). Grand-mean intensity normalization of the entire 4D dataset was also carried out by a single multiplicative factor; highpass temporal filtering (Gaussian-weighted least-squares straight line fitting, with sigma=45.0s). Volumes were brain extracted using FSL BET [44] and the medial functional volume was coregistered to each participant’s MPRAGE image (df = 6) and the T1 MNI standard template (df = 12) using a rigid-body transformation in FMRIB’s Linear Image Registration Tool (FLIRT) [43,45]. Time-series statistical analysis was carried out using FILM with local autocorrelation correction [46].

## Regions of interest

As noted earlier, our hypotheses centered on the roles of the AT and PM networks in using the higher-order conceptual structure of the task to guide learning. To address this question, we used thirty-six regions of interest (ROIs) within the PM and AT networks (18 AT ROIs, 18 PM ROIs). PMAT ROIs were defined as 6 mm spheres centered on coordinates identified in an independent dataset on the basis of a comparison of PHc and PRc resting-state functional connectivity (RSFC) [47]. Spheres were non-overlapping and spatially separated by a distance of at least 12 mm. PMAT network assignment of each sphere was determined on the basis of resting state networks identified in a separate independent dataset [48]. To determine network assignment in this independent dataset, Ritchey et al. applied a community detection algorithm to RSFC time courses extracted from the spheres identified based on functional connectivity [47], allowing for an identification of spheres that showed greater within vs. between network connectivity [48]. All ROIs were transformed into standard MNI space using FSL FLIRT with nearest-neighbor interpolation.

## FMRI statistical analysis

We conducted three fMRI analyses to assess the involvement of the PM and AT networks in indexing dissociable aspects of learning in a task with a higher-order conceptual structure. To assess network involvement in the conceptual integration of cue pairs with shared feature relationships (“Cue-based integration”) and incremental updating of the conceptual cluster space (“Feedback-based updating”), we conducted a computational model-based fMRI analysis. In order to rule out whether the results of the model-based analyses could be attributed to task performance alone, we also conducted a separate accuracy-based univariate analysis assessing PM and AT network activation during the cue pair and outcome periods of correct relative to incorrect trials. Finally, in order to provide evidence supporting dissociable PM and AT networks in the current task, we conducted an activation profile similarity analysis using estimates from both the model- and accuracy-based analyses.

## Computational model-based fMRI analysis

The RMC was individually fit to each participant’s learning data to generate trial-by-trial estimates of Cue-based integration and Feedback-based updating (see section on *Model-based analyses*). Task activation was assessed using a parametric univariate analysis implemented in FSL. Individual modulated and unmodulated regressors were constructed for each iteration through the full set of Cue 1 – Cue 2 – Outcome associations (three iterations per run). Model-derived estimates were modeled on an iteration-by-iteration basis to avoid confounding model-based learning measures with effects related to time and other random variance. To estimate neural activation associated with the conceptual integration of cue pairs with shared feature relationships, three model-derived Cue-based integration parametric regressors (one for each iteration) were included in the first-level GLM. Parametric regressors for Cue-based integration were modeled at the onset of Cue 1 with a duration of 4.5 seconds to include the full Cue 1 and Cue 2 presentation period. To estimate neural activation associated with the updating of the conceptual cluster space in response to feedback information (“Feedback-based updating”), three model-based parametric regressors reflecting Feedback-based updating were also included. Each Feedback-based updating parametric regressor was modeled at the onset of the Feedback period with a duration of 2 seconds. Cue and Feedback-based parametric regressors were mean-corrected separately on an iteration-by-iteration basis. To model the mean activation associated with the Cue period, three 4.5 second unmodulated boxcars (one regressor for each iteration) were modeled at the onset of Cue 1 with a duration of 4.5 seconds. Mean activation associated with the Feedback period was modeled similarly, with three separate 2 second unmodulated boxcars (one regressor for each iteration) beginning at the onset of Feedback. The first iteration of the first learning run was modeled as a 7 second unmodulated nuisance regressor to allow for stabilization of model estimates following one complete iteration through the full set of cue-pair outcome associations. Regressors were convolved with a double-gamma hemodynamic response function prior to model estimation.

In order to assess the parametric effects of Cue-based integration and Feedback-based updating in each run, first level contrasts for Cue-based integration and Feedback-based updating were computed separately. Contrasts included regressors from all iterations. To average contrast estimates over all six learning runs, a second-level fixed effects model was carried out by forcing the random effects variance to zero in FLAME (FMRIB’s Local Analysis of Mixed Effects) [49–51]. Participant-level statistical contrast maps for Cue-based integration and Feedback-based updating were transformed to standard MNI template space for subsequent analysis.

To characterize the roles of the PM and AT networks in Cue-based integration and Feedback-based updating, mean parameter estimates were extracted for each ROI from each contrast and averaged within network, yielding four estimates per participant (AT network – Cue-based integration, PM network – Cue-based integration, AT network – Feedback-based updating, PM network – Feedback-based updating). Contrast estimates were submitted to a repeated-measures ANOVA assessing the effects of Network (PM, AT) and Trial period (cue, feedback), and followed up with planned two-tailed paired comparisons.

## Accuracy-based fMRI analysis

To rule out whether parametric activation associated with Cue-based integration and Feedback-based updating was merely a reflection of task accuracy, we conducted a separate accuracy-based analysis to assess cue and feedback activation during trials with correct relative to incorrect category outcome decisions. In order to ensure sufficient trial numbers to estimate contrasts for correct and incorrect trials, functional data from runs one through three were concatenated into an “Early learning” period, and functional data from runs three through six were concatenated to create a “Late learning” period. First-level modeling of each learning period included four regressors of interest for the cue and outcome periods of trials associated with correct and incorrect category outcome decisions. Additionally, two unconvolved nuisance regressors were included to model the effect of run. To assess cue and feedback-based activation for correct > incorrect trials across early and late learning, a second-level fixed-effects model was carried out by forcing the random effects variance to zero in FLAME [49–51]. Participant-level statistical contrast maps for Cue: correct > incorrect and Feedback: correct > incorrect were transformed to standard MNI template space for subsequent analysis. Subject-level contrast estimates were extracted for each ROI from each contrast and averaged, yielding four estimates per network per participant (AT network – Cue: correct > incorrect; PM network – Cue: correct > incorrect; AT network – Feedback: correct > incorrect; PM network – Feedback: correct > incorrect). Contrast estimates were submitted to a repeated-measures ANOVA assessing the effects of Network (PM, AT) and Trial period (cue, feedback), and followed up with two-tailed t-tests to assess planned paired comparisons.

## Activation profile similarity analysis

In order to validate our analyses on data from the AT and PM networks, we tested the appropriateness of grouping individual ROIs according to this network framework. Following Ritchey et al. [48] we ran an “activation profile similarity analysis,” to test whether regions in the same putative network showed more similar profiles of activation across the different experimental conditions than did regions in different networks. Mean subject-level contrast estimates from each condition of interest (Cue-based integration; Feedback-based updating; Cue: correct > incorrect; Feedback: correct > incorrect) were z-transformed and extracted from each ROI. This procedure yielded an activation matrix with a separate four-element activation vector for each ROI. To measure the similarity of activation profiles across all pairs of ROIs, the activation matrix was correlated using Pearson’s *r* and sorted by network affiliation. In order to assess functional homogeneity within network, the resulting activation similarity correlation matrix was compared to a model matrix, where pairs of ROIs within the same network were represented with a value of 1 (perfect correlation) and ROI pairs in different networks were represented with a value of 0 (no correlation). Both the activation similarity correlation matrix and model matrix were vectorized and compared with Kendall’s Tau, a non-parametric measure of statistical dependence. Additionally, activation similarity correlation values were averaged across regions within the PM and AT networks (within-network) and compared to the average across regions in the PM and AT networks (between-network). Correlation values were Fisher z-transformed prior to comparison via a two-tailed paired t-test.

## Results

### Behavioral results and computational model fits

Behavioral performance indicated that participants were, on average, able to learn associations between pairs of cues and outcomes across six learning runs (Fig 3). When submitting average performance across runs to a repeated-measures ANOVA, there was a significant main effect of Run [F_5,120_ = 39.5, p <. 000001], indicating that outcome decisions improved significantly across the six task runs. Consistent with this idea, outcome decisions were not significantly different from chance (50%) until the fourth (p < 0.05, binomial test), fifth (p <. 0001, binomial test) and sixth (p <. 0001, binomial test) task runs. Additionally, outcome decision reaction times were found to decrease significantly across runs [F_5,120_ = 2.82, p <. 05]. Despite a group-level improvement in performance across learning runs, there was a large amount of variability in individual participant ability to learn Cue 1 – Cue 2 – Outcome associations (Fig 3). As such, the standard deviation of the group accuracy score (percent correct) increased from 0.06 in the first run to. 18 on the final learning run.

**Fig 3.**
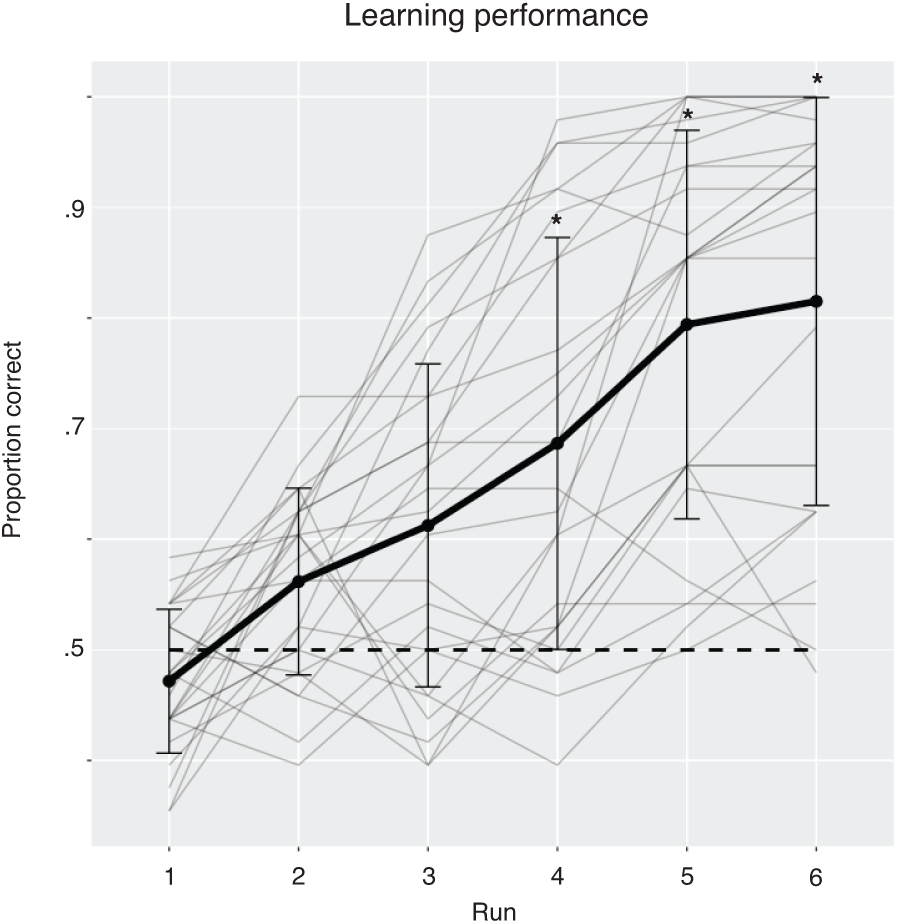
Learning performance. On average, participants gradually learned to choose outcomes correctly across six runs of scanned learning, however there was a large amount of variability in individual participant performance. Chance performance (50%) is plotted by a dashed line. Mean subject performance plotted in black. Individual subject learning curves plotted in grey. Error bars reflect +/- 1 standard deviation from the mean. * p <. 05, binomial test.

Next, we assessed whether individual participant behavioral estimates from the RMC were able to approximate subject performance during the learning task (Fig 4). On average, the R^2^ across all participants was. 47 with a standard error of +/- 0.0559, suggesting that the RMC provides a good fit to the observed behavioral data despite the large amount of variability observed across participants.

**Fig 4.**
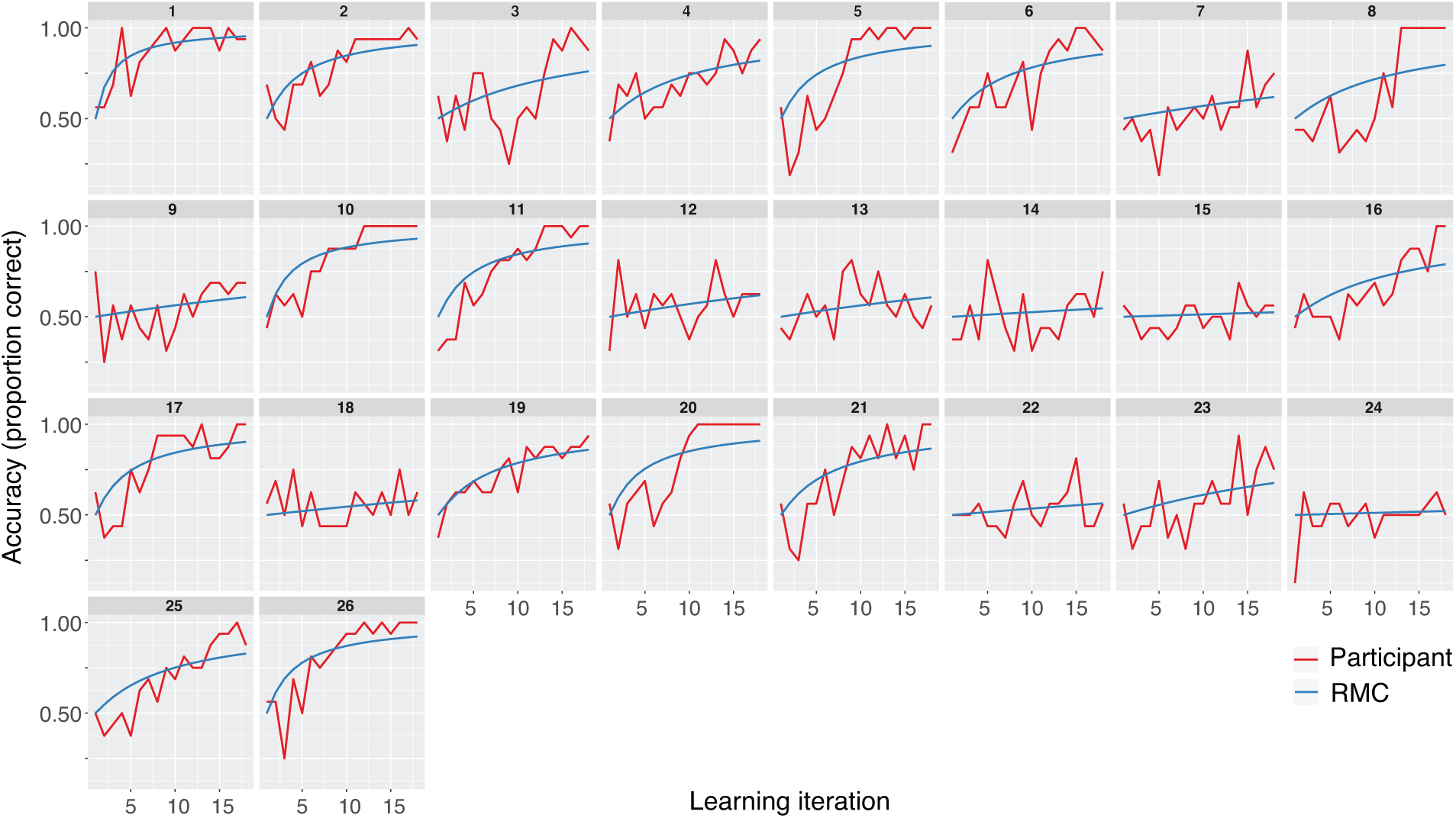
Individual participant behavioral model fits.

## Linking of experiences in memory is associated with facilitated learning

Although the task consisted of sixteen individual cue-pair outcome associations, capitalizing on shared feature relationships could reduce the learning problem and facilitate correct outcome decisions. To assess evidence in support of this idea, we turned to the coupling probability, or a single model-derived value that describes each participant’s tendency to link cue pair-outcome associations (see section on *Model-based analysis*). We observed a significant positive correlation between the coupling probability and the slope of each participant’s learning curve. Specifically, larger coupling probabilities were associated with faster learning rates in the task (Fig 5, [r(24) =. 558, p =. 003]).

**Fig 5.**
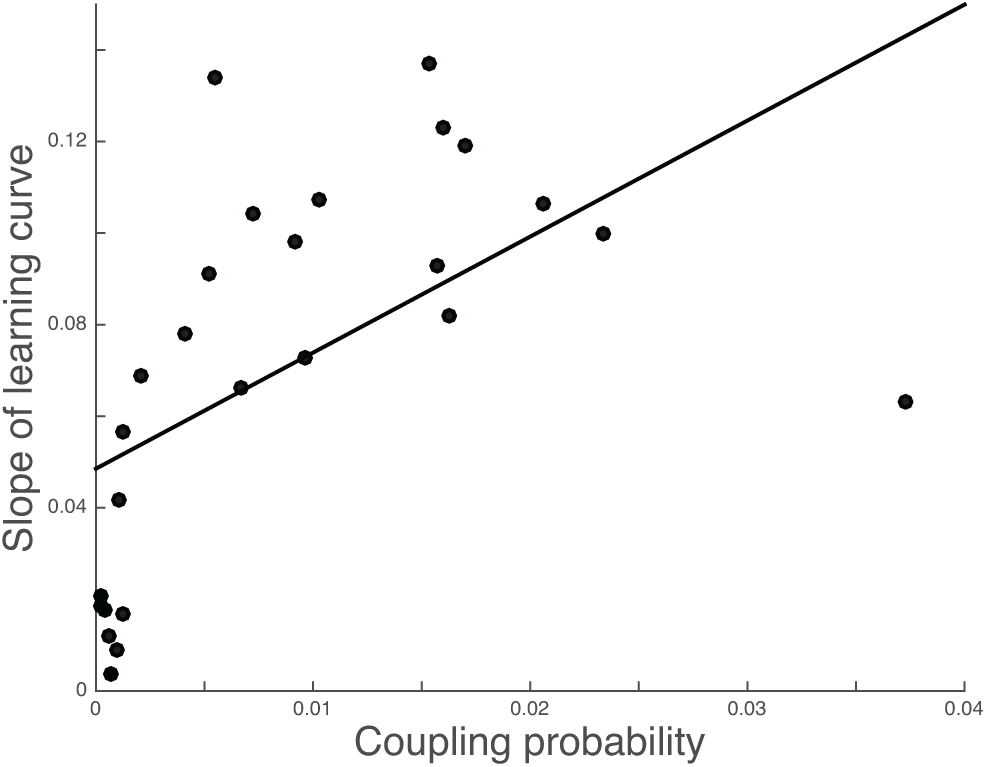
Larger coupling probabilities are associated with faster learning rates. Individual subject coupling probabilities, or a model-derived metric where higher values reflect a stronger tendency to link information in memory, is positively associated with the rate of learning.

## Evidence for dissociable PM and AT networks

As noted earlier, prior evidence is consistent with the idea that regions in the AT or PM networks (or both) could contribute to dissociable aspects of learning conceptual information in this task. Prior work suggests that there is substantial similarity in the extent to which regions within the same network are recruited during different task conditions [48]. To conceptually replicate these results and to verify the appropriateness of grouping ROIs according to the PMAT framework [13], we conducted an activation profile similarity analysis [48] in which we quantified the similarity of task-based activation profiles across regions within and across each network (Fig 6). If individual ROIs are operating in concert with other within-network regions and processing similar types of information in the learning task, we should see similar activation values across regions that belong to the same network. Additionally, this relationship should be true for activation values derived from all available contrasts.

**Fig 6.**
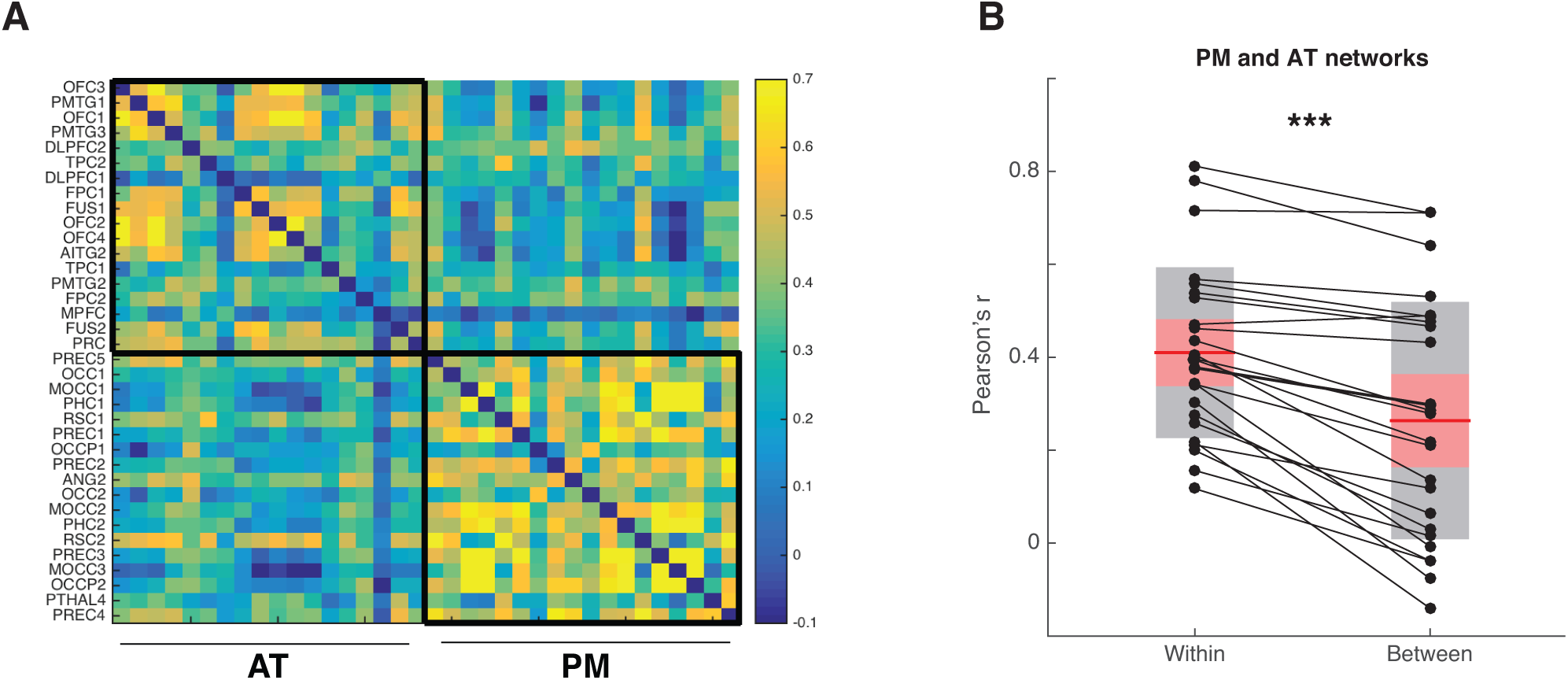
Activation profile similarity analysis. Regions within the PM and AT networks show high within but not between network activation profile similarity. (A) Activation profile similarity values were assessed by correlating mean z-transformed contrast values from each ROI extracted from each of the four contrasts of interest (Model – Cue-based prediction, Model – Feedback-based updating, Accuracy – Cue: Correct > Incorrect, Accuracy – Outcome: Correct > Incorrect). Higher correlation values indicate that a pair of ROIs displayed a more similar pattern of activation across contrasts. (B) Activation profile similarity correlations were significantly higher between ROIs that were from within the same network relative to across different networks. Grey-shaded box denotes standard deviation. Red shaded box denotes 95% confidence interval. Individual participant activation profile similarity values plotted in black. *** p <. 0001

Activation profile similarity scores are computed by correlating univariate activation vectors across all pairs of ROIs within and across networks. A pair of ROIs will display high correlation, or high activation profile similarity, if both regions exhibit a comparable pattern of relative activation or deactivation across the four contrasts derived from the computational model- and accuracy-based analyses (Model – Cue-based integration; Model – Feedback-based updating; Accuracy – Cue: correct > incorrect; Accuracy – Feedback: correct > incorrect). To assess whether regions within each network display similar activation profiles, correlation values were sorted by network affiliation, vectorized, and compared with an idealized model matrix, where correlations between regions within the same network were represented as 1 (perfect correlation) and correlations for ROI pairs in different networks were represented as 0 (no correlation). The Kendall’s tau correlation between activation profile similarity values and the model matrix was statistically significant (Kendall’s tau =. 323, p <. 0001), suggesting that activation profile similarity was higher across pairs of ROIs within the same network relative to ROI pairs across networks. A complimentary analysis directly comparing average within-relative to between-network correlations was consistent with these results, demonstrating significantly higher correlation values across pairs of ROIs within the same network relative to across networks [t(24) = 8.11, p <. 0001]. Indeed, nearly every participant demonstrated higher within-relative to between-network activation profile similarity values, providing further evidence that regions in the PM and AT networks are engaged in separable processes as participants completed the task. Having established the validity of the distinction between the AT and PM networks, our next analyses focused on characterizing how these networks contributed to the development and use of conceptual information by relating them to two key indices from the computational model – Cue-based integration and Feedback-based updating.

## Differential PM and AT network involvement in supporting Cue-based integration and Feedback-based updating

As noted in the introduction, there is good reason to think that the AT or PM networks, or both, would contribute to concept acquisition in this task. Based on previous work implicating the AT network in supporting information about the meaning of objects and complex conjunctions of features, we hypothesized that regions in this network should collectively track conceptual knowledge reflecting shared feature relationships. Alternatively, because shared feature relationships are built on the local context of each trial, one might expect regions in the PM network to preferentially represent Cue-based integration. A third possibility is that the AT and PM networks might play complementary roles in Cue-based integration. Building on recent proposals that the PM network supports representations of relevant relationships in the environment, we also hypothesized that this network would track trial-by-trial updates to the conceptual cluster space.

To test these hypotheses, parameter estimates indexing activation related to Cue-based integration and Feedback-based updating were submitted to a repeated measures ANOVA with factors for Network (PM, AT) and Trial period (cue, outcome). Results revealed a significant main effect of Network (F_1,24_ = 9.89, p <. 01) and a significant Trial period by Network interaction (F_1,24_ = 7.34, p <. 012) (Fig 7, left panel). No main effect of Trial period was observed (F_1,24_ = 0.009, p =. 92). Follow-up paired comparisons revealed that Cue-based integration estimates were significantly lower in the AT network relative to the PM network [t(24) = 3.22, p <. 01]. Additionally, one sample t-tests revealed that Cue-based integration parameter estimates were significantly different from zero in the AT network [t(24) = −2.9, p <. 01] but were not significantly different from zero in the PM network [t(24) = 1.36, p = 0.18] (Fig 7A). These results suggest that activity in the AT network reflected the integration of cue pair information into existing concepts, consistent with the use of the higher-order conceptual structure of the task (see S1 Fig for results from individual PM and AT network ROIs). The PM network, on the other hand, did not display evidence for involvement in Cue-based integration.

**Fig 7.**
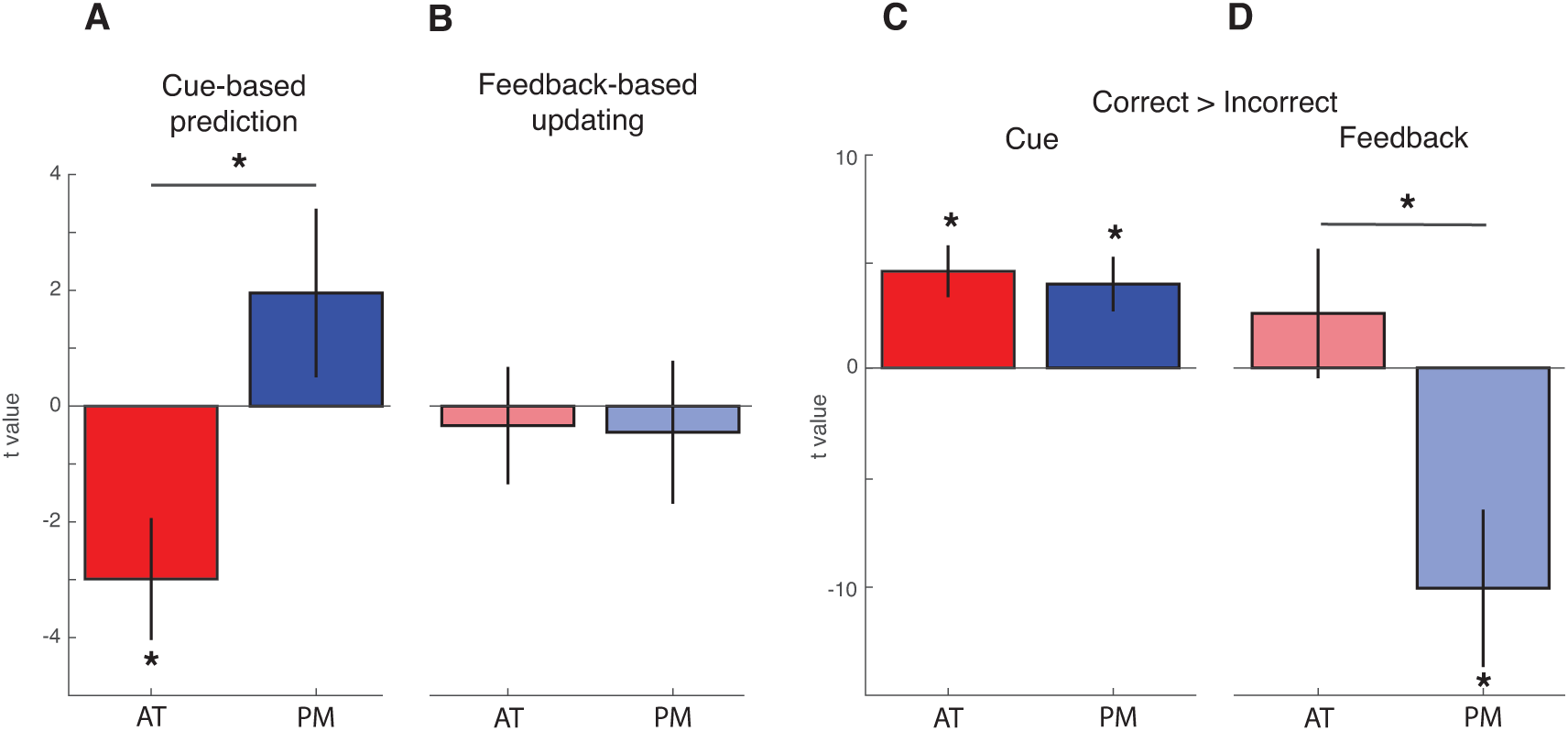
Model- and accuracy-based analyses in the PM and AT networks. (A) AT network is involved integrating cue pairs within an existing cluster. Values above zero denote greater probability of assigning cue pairs to a novel cluster. Values below zero denote greater probability of capitalizing on shared features to assign cue pairs to an existing cluster. (B) Parametric activation reflecting feedback-based updating, or incremental changes to the conceptual cluster space following feedback. Positive values reflect more updating. Values below zero denote less updating. (C) Accuracy-based univariate analyses reveal that both networks demonstrate greater activation for cue pairs associated with subsequent correct relative to incorrect predictions. (D) Outcome-related univariate activation in the PM network is significantly greater for incorrect relative to correct predictions. Error bars denote standard error of the mean. * p <. 05

Results also did not reveal support for either PM or AT network involvement in indexing Feedback-based updating. Neither PM network nor AT network estimates were significantly different from zero [PM: t(24) = −0.37, p =. 71; AT: t(24) = -.341, p =. 73], and estimates did not differ significantly across networks [t(24) = 0.15, p = 0.88].

## PM and AT networks jointly support accuracy during learning

In order to determine whether computational model-based fMRI analyses were able to provide unique insights into the involvement of the PM and AT networks in the updating and use of conceptual representations, we also assessed the involvement of these networks in a univariate accuracy analysis (see S2 Fig for results from individual ROIs). One sample t-tests revealed that parameter estimates were reliably different from zero in the AT network during the Cue period only [t(24) = 3.94, p <. 001], whereas PM network estimates were significantly different from zero during both the Cue [t(24) = 3.24, p <. 01] and Outcome [t(24) = −2.85, p <. 01] trial periods (Fig 7, right panel). The resulting average parameter estimates were entered into a repeated measures ANOVA with factors for Network (PM, AT) and Trial period (Cue, Outcome). Results revealed significant main effects of Trial period (F_1,24_ = 5.38, p <. 05), Network (F_1,24_ = 27.1, p <. 00001) and a significant Trial period by Network interaction (F_1,24_ = 18.9, p <. 0001). Follow-up paired comparisons revealed a significant difference in feedback activation for the PM and AT networks [t(24) = 5.4, p <. 0001, Fig 7B] with no differences across networks during the Cue period [t(24) =. 46, p =. 64]. These findings indicate that although the PM and AT networks differentially contributed to feedback learning, both displayed significant Cue period activity during trials that were associated with correct outcome decisions. These results are in contrast to the model-based analysis, which show differential PM and AT activity during the cue period.

## Discussion

Although recent neuroimaging investigations have focused on elucidating brain regions involved in concepts formed on the basis of simple features, concepts can also reflect higher-order relationships, including shared features across entities in the environment. Here, we used computational modeling-based fMRI to assess how these types of conceptual representations are constructed and used during learning. Behavioral and model results revealed that participants who showed a tendency to link information in memory, as evidenced by the fitted coupling probability value, also had faster learning rates, suggesting that use of the conceptual structure of the task facilitated learning. Analyses of fMRI data revealed that activity in the AT network tracked the integration of cue pair information into existing conceptual clusters (“Cue-based integration”), consistent with proposals that this network encodes information about the meaning or significance of objects [13].

The present study stressed learning to integrate cues that, in isolation, were not diagnostic, in order to predict future outcomes. Additionally, although one could learn the simple associations between cue pairs and outcomes, performance could be optimized by learning concepts reflecting related events. According to cluster-based models of categorization, such as the RMC [1], categories are indirectly represented by grouping past experiences into conceptual clusters that include information about items and the categories that they had been associated with. We were especially interested in understanding how the AT and PM networks might support this process given evidence indicating that regions in the AT network represent the conceptual features of objects [20,52–55] and work that has found representations of contextual information in the PM network [25,27,32].

Results derived from the model-based fMRI analysis provide novel evidence that the AT network is involved in the integration of cue pair information into concepts that reflect shared features (“Cue-based integration”). These results complement existing fMRI evidence that have largely taken a region-specific approach as opposed to an investigation of network-wide effects. For example, a number of investigations have found evidence that the PRc is sensitive to conceptual processing [56–59] and complex conjunctions of stimulus features [60–65]. Extending these results, PRc has also been implicated in representing the significance or meaning of stimuli [17,20,21,52,66]. Consistent with a role for the AT network in integrating cue information into existing conceptual representations, prior work has also found that the OFC is critically involved in integrating prior experience with current evidence to support decisions [18,23,67,68]. Research assessing OFC involvement in learning has also found evidence that this area represents task-relevant information that cannot be directly observed [69–75], similar to the higher-order conceptual structure of the current task. Convergent results from neuropsychology have also found impairments in semantic memory and multimodal semantic processing in patients with damage to the anterior temporal lobes following semantic dementia, herpes simplex encephalitis, or temporal lobe resection [76–80], consistent with the idea that regions in this network represent conceptual and meaning-based information.

Interestingly, we did not find evidence for PM network involvement in indexing Cue-based integration, despite the importance of local context. One possible explanation for this is the type of stimuli used in the task. Regions in the PM network, including PHc and RSc, have typically been associated with learning and memory of contextual information that is principally spatial or location-based [32,81–87]. The AT network, on the other hand, has been found to be particularly sensitive to object stimuli [64,65]. It is possible that the PM network might engage in Cue-based integration in a similar task using scene or spatial images rather than objects.

We additionally investigated PM involvement in tracking trial-by-trial updates to the conceptual cluster space (“Feedback-based updating”) given recent proposals that this network represents the full set of relationships that are relevant in the environment [13]. Results did not reveal evidence to suggest that PM network activity tracked Feedback-based updating, although PM network activation was significantly higher on incorrect trials relative to correct trials. One interpretation of these results is that the learning problem in the current task did not encourage large-scale reorganizations of the conceptual cluster space after trial-by-trial feedback [42,88]. This interpretation is consistent with individual participant behavioral learning curves and model fits, which largely depict a gradual increase in performance across the six learning runs (Fig 4). Future work will be required to assess whether and how the PM network is involved in representing or updating the full set of relationships active in the current environment.

It is also interesting to consider how the current investigation complements and extends previous work. In particular, recent model-based fMRI investigations have evaluated neural evidence for different forms of category learning [89], assessed the brain areas involved in the reorganization of conceptual knowledge following changes in attention and goals [41], and identified brain regions involved in category learning with stimuli that are consistent and inconsistent with category rules [5,6]. Together these investigations have largely found evidence for hippocampal and striatal involvement, and recent theoretical work has suggested that the hippocampus may play a key role in concept formation and organization [90]. These investigations, however, have largely defined conceptual representations on the basis of simple perceptual features or associations, leaving open the question of how concepts defined on shared feature relationships are developed and used to guide learning. The current investigation suggests that these types of higher-order concepts are preferentially represented by the AT network, consistent with this network’s role in representing the meaning of objects in the environment and information about complex feature conjunctions.

## Acknowledgements

The authors would like to thank Maureen Ritchey for helpful discussion.

## Supporting information

**S1 Fig.**
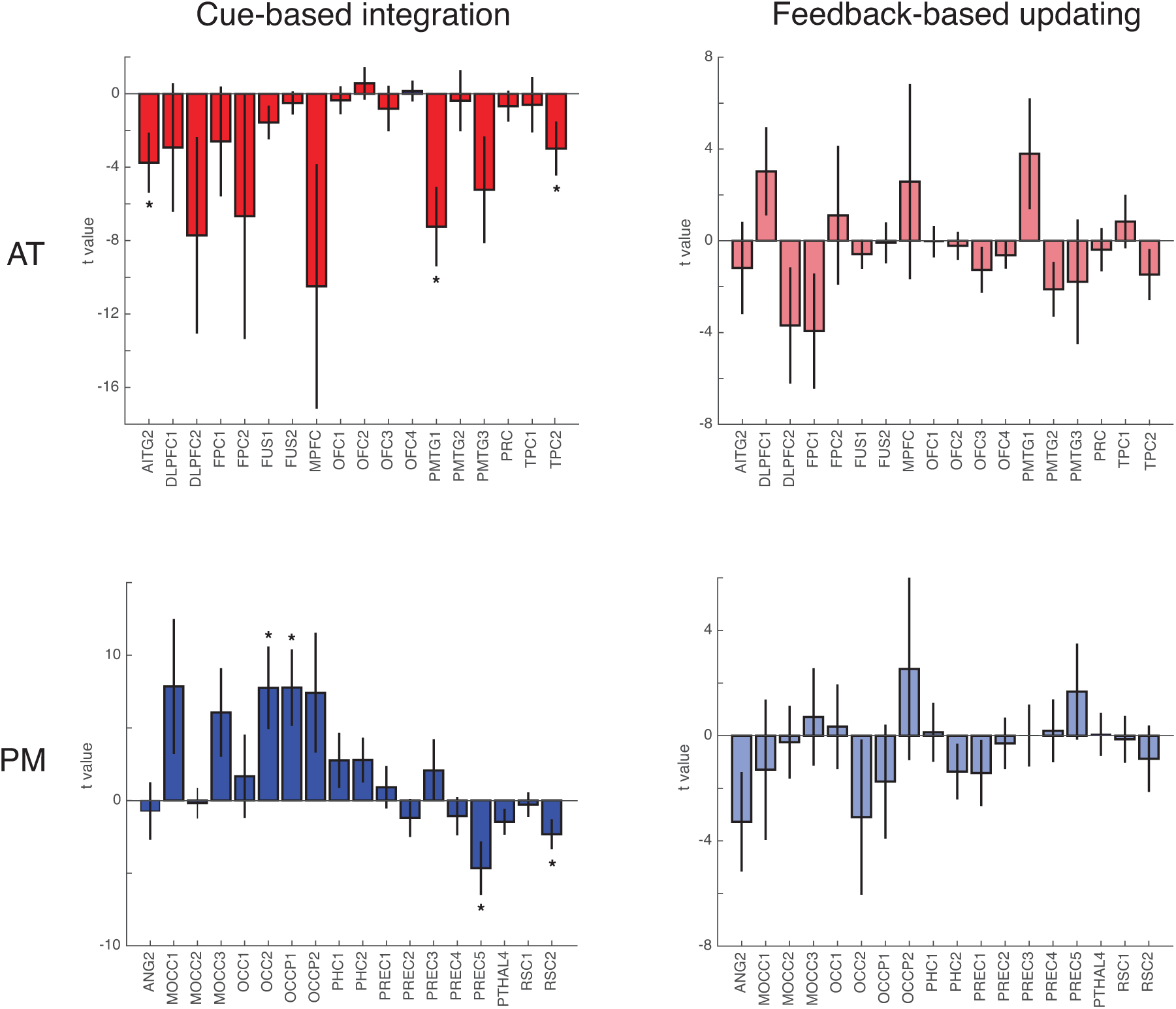
Model-based results for individual regions in the PM and AT networks. Error bars denote standard error of the mean; * p <. 05, one sample t-test.

**S2 Fig.**
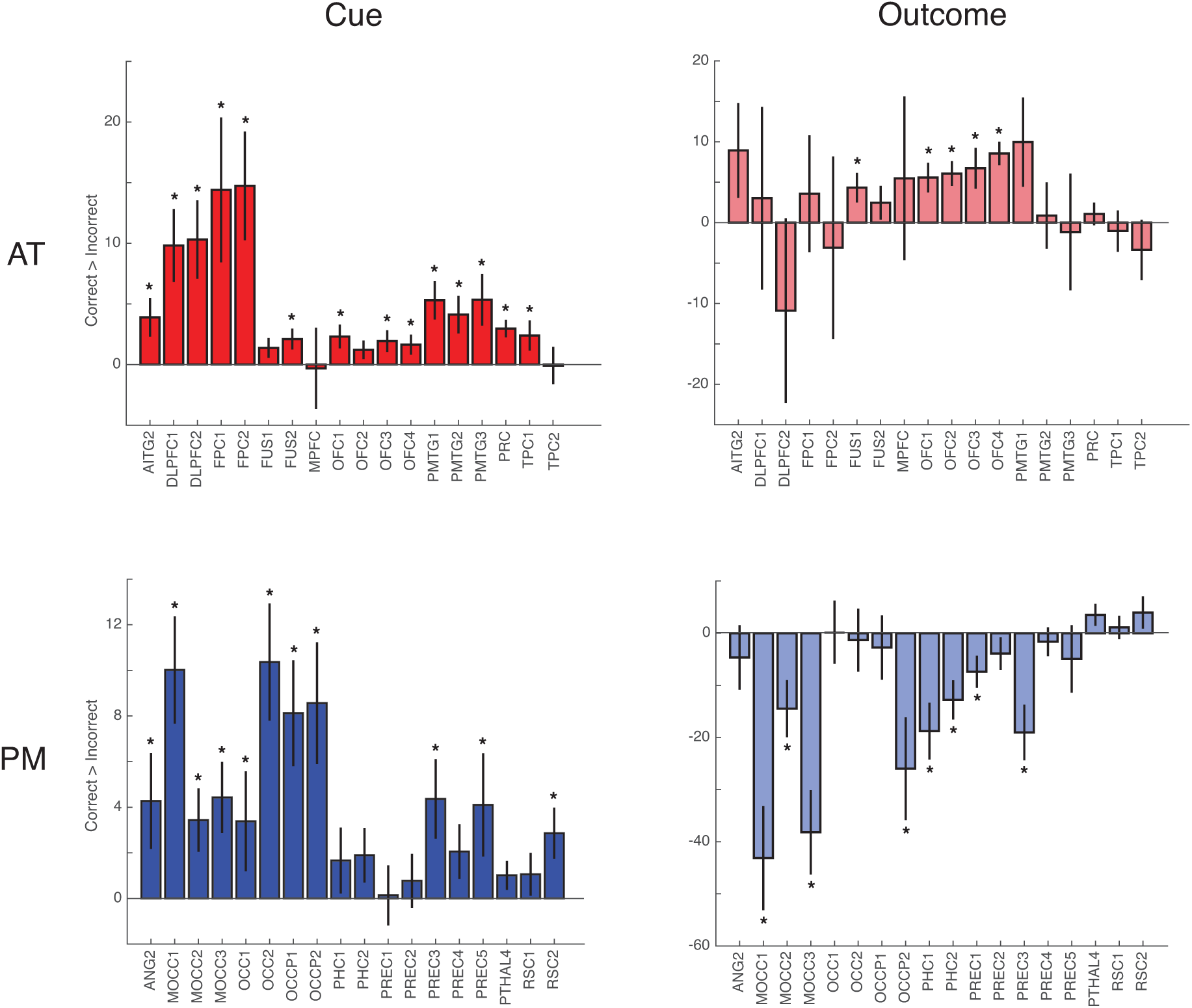
Accuracy-based results for individual regions in the PM and AT networks. Correct > Incorrect contrast results. Error bars denote standard error of the mean; * p <. 05, one sample t-test.

